# ReorientExpress: Reference-free orientation of nanopore cDNA reads with deep learning

**DOI:** 10.1101/553321

**Authors:** Angel Ruiz-Reche, Akanksha Srivastava, Joel A. Indi, Ivan de la Rubia, Eduardo Eyras

**Affiliations:** Pompeu Fabra University, E08003, Barcelona, Spain; EMBL Australia, The John Curtin School of Medical, Australian National University, Acton ACT 2601, Canberra, Australia; Instituto de Medicina Molecular, Faculdade de Medicina, Universidade de Lisboa, Lisbon, Portugal; Hospital del Mar Research Institute, E08003, Barcelona, Spain

## Abstract

We describe ReorientExpress (https://github.com/comprna/reorientexpress), a method to perform reference-free orientation of transcriptomic long sequencing reads. ReorientExpress uses deep-learning to correctly predict the orientation of the majority of reads, and in particular when trained on a closely related species or in combination with read clustering. ReorientExpress enables long-read transcriptomics in non-model organisms and samples without a genome reference and without using additional technologies.

## Background

Long-read sequencing technologies allow the systematic interrogation of transcriptomes from any species. However, functional characterization requires knowledge of the correct 5’-to-3’ orientation of reads. Oxford Nanopore Technologies (ONT) allows the direct measurement of RNA molecules in the native orientation (Garalde et al., 2018), but the sequencing of complementary-DNA (cDNA) libraries yields generally a larger number of reads (Garalde et al., 2018; Workman et al., 2018). Although strand-specific adapters can be used, error rates hinder their correct detection. Current methods to analyze nanopore transcriptomic reads rely on the comparison to a genome or transcriptome reference (Workman et al., 2018; Wyman and Mortazavi, 2019) or on the use of additional technologies such as in ‘hybrid sequencing’, which employs long and short read data (Fu et al., 2018), which limits the applicability of rapid and cost-effective long-read sequencing for transcriptomics beyond model species. To facilitate the *de novo* interrogation of transcriptomes in species or samples for which a genome or transcriptome reference is not available, we have developed ReorientExpress, a new tool to perform reference-free orientation of ONT reads from a cDNA library. ReorientExpress uses deep neural networks (DNNs) to predict the orientation of cDNA long-reads independently of adapters and without using a reference. ReorientExpress predicts correctly the orientation of the majority of cDNA reads, and in particular when trained on a related species or in combination with read clustering, thereby enabling the reference-free characterization of transcriptomes.

## Results

### Sequence-based prediction of read orientation

ReorientExpress approach builds on the hypothesis that RNA molecules present sequence biases and motifs relevant to their metabolism, generally related to protein binding sites (Hentze et al., 2018; Rissland, 2017). Despite potential sequencing errors, these signals may still be largely present in a long read and therefore enable the identification of the right orientation of a cDNA read. ReorientExpress implements two types of Deep Neural Network (DNN) models to classify long reads as being in the forward (5’-to-3’) orientation or in the reverse-complement-orientation (Fig. 1a and b). The first DNN model is a multilayer perceptron (MLP) with 5 hidden layers, the last layer providing the probability that a read is not in the correct orientation, and with dropout layers to reduce overfitting (Additional file 1: Table 1). In this model, an input sequence is represented in terms of the frequency of short motifs (k-mers for k=1,...,5). This ensures a fixed-size input for molecules of different lengths and accounts for the fact that sequencing errors may not allow to correctly capture longer sequence patterns. Furthermore, neural networks work better with input values between 0 and 1. Accordingly, the input of the MLP model is a matrix of normalized k-mer counts from the input sequences, with k from 1 to any length specified as parameter (default=5) as shown in Fig. 1a. Although MLPs are simpler and faster to train and run, they do not capture the dependencies with sequence context like Convolutional Neural Networks (CNNs). For this reason, ReorientExpress also implements a CNN as an alternative model with a similar architecture to lenet (Lecun et al., 1998) with 3 convolutional layers, 3 pooling layers and 2 dense layers, with different filter sizes (Additional file 1: Table 2). As input for the CNN model, sequences were divided into overlapping windows of fixed size, which were then transformed using one-hot encoding (Methods). The orientation of a read is based on the mean of the posterior probabilities for all windows in a read for forward and reverse orientation and thereafter by selecting the one with the highest value (Fig 1b). Regardless of the DNN model, ReorientExpress can be trained from any set of sequences with known orientations, like a transcriptome annotation or ONT direct RNA sequencing (DRS) reads (Methods).

**Figure 1.**
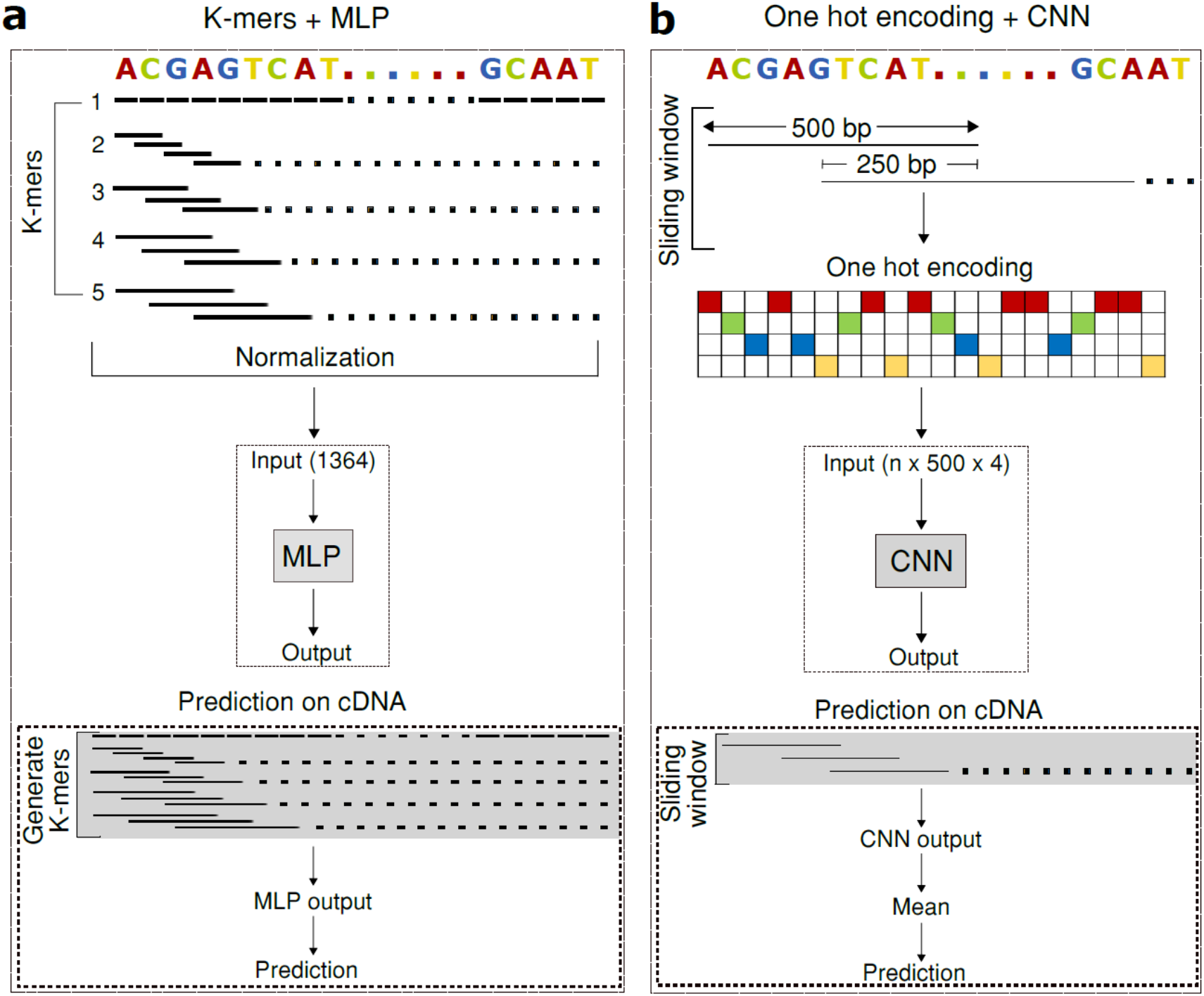
ReorientExpress Deep Learning models. ReorientExpress implements two deep neural networks (DNNs) to predict the orientation of cDNA long reads. **(a)** A multilayer perceptron (MLP) is trained on k-mer frequencies extracted from sequences of known orientation. For each test read, the orientation is predicted using the k-mer frequencies of the read as input. **(b)** A convolutional neural network (CNN) is trained on 500nt sliding windows from sequences of known orientation, using one-hot encoding for each window (Methods). Prediction is performed by scoring all windows in a test read and calculating the mean score independently for each orientation.

To test ReorientExpress, we first trained the MLP model on 50,000 random transcripts from the human annotation using k-mers from k=1 to k=5. To enable the accuracy evaluation with ONT cDNA reads with unknown orientation, we first mapped human ONT cDNA reads to their respective transcriptomes using minimap2 (Li, 2018). Thereafter, we selected only uniquely mapped reads with maximum quality (MAPQ=60), and assigned to each read the strand from the matched annotated transcript (Methods). Using these orientations as ground truth, the MLP model trained on human transcriptome yielded an average precision of 0.84 and a recall of 0.83 in human cDNA reads (Fig. 2a) (Additional file 1: Table 3). The CNN model trained on the human transcriptome showed slightly better results on the human cDNA reads (Fig. 2a) (Additional file 1: Table 3). We proceeded in a similar way with *S. cerevisiae* cDNA reads (Garalde et al., 2018), as described above for human cDNA reads. ReorientExpress trained on the *S. cerevisiae* annotation (Methods) yielded an average precision and recall of 0.93 in ONT cDNA reads from *S. cerevisiae* (Fig 2b) (Additional file 1: Table 3). Similar to the above observation in human cDNA, the accuracy of the CNN model on the *S. cerevisiae* reads was slightly better than the MLP model (Fig 2b) (Additional file 1: Table 3). Application of the MLP or CNN models trained on human transcriptome to direct RNA sequencing (DRS) reads yielded better accuracy for human DRS reads compared to human cDNA reads. However, the accuracy decreased for both MLP and CNN model trained on *S. cerevisiae* transcriptome and tested on *S. cerevisiae* DRS reads compared to *S. cerevisiae* cDNA reads (Additional file 1: Table 3).

**Figure 2.**
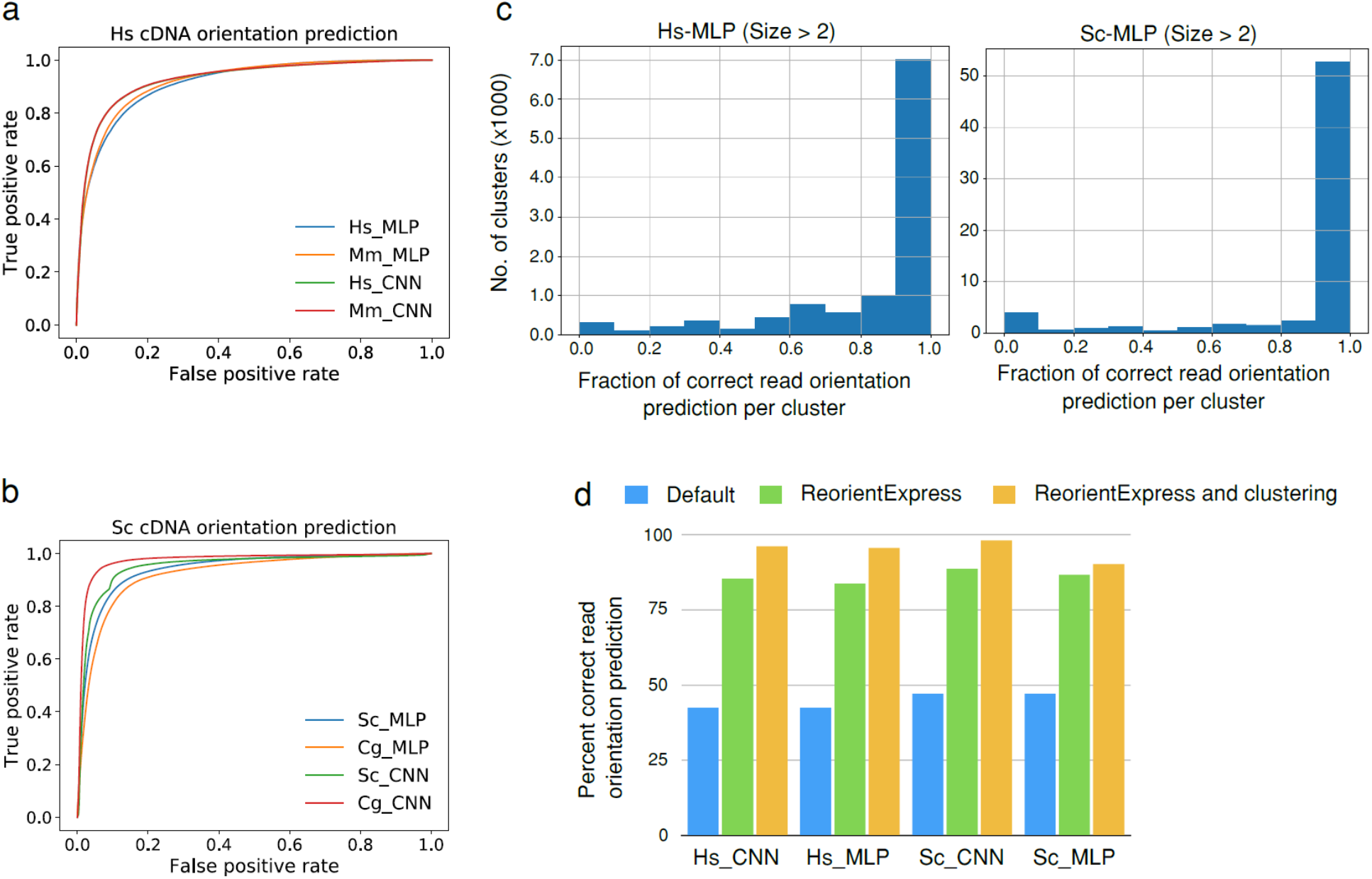
ReorientExpress accuracy analysis. **(a)** Receiving Operating Characteristic (ROC) curves, representing the false positive rate (x axis) versus the true positive rate (y axis) for the prediction of the orientation of human ONT cDNA reads with the multilayer perceptron (MLP) and convolutional neural network (CNN) models trained on either the human (Hs) or the mouse (Mm) transcripts. **(b)** ROC curves for the prediction of the orientation of yeast ONT cDNA reads with the MLP and CNN models trained on either the *S. cerevisiae* (Sc) or *C. glabrata* (Cg) transcripts. **(c)** Number of clusters (y axis) according to the proportion of human ONT cDNA reads in the cluster with orientation correctly predicted by ReorientExpress (x axis) with the MLP model trained on the human transcriptome (left panel) (Hs-MLP) or the *S. cerevisiae* transcriptome (right panel) (Sc-MLP). We show clusters with more than 2 reads. Similar plots for all clusters (>1 read) and for the CNN model are given in Additional file 1: Figure 1. **(d)** Comparison of the proportion of human (Hs) or *S. cerevisiae* (Sc) cDNA reads correctly oriented in three cases: taking the default orientation from the FASTQ file (Default) in blue, using the CNN and MLP ReorientExpress models in green, and using a majority vote in clusters to predict the orientation of all reads in each cluster (ReorientExpress and clustering) in orange. Clustering and predictions in (c) and (d) were performed with all labelled cDNA reads (Methods). Models used to on the total set of labelled cDNA reads in this figure were trained on 50,000 randomly selected transcript sequences from the annotation, or all of them if there were less (*S. cerevisiae* and *C. glabrata*).

To demonstrate the suitability of ReorientExpress to predict the orientation of cDNA reads from samples without a genome or transcriptome reference available, we mimicked this situation by building DNN models in one species and testing them on a related species. We thus trained an MLP model (k=1,…,5) with the mouse transcriptome. This model tested on human ONT cDNA reads showed a precision of 0.79 and recall of 0.71, which is comparable to the MLP model trained on human data (Fig. 2a) (Additional file 1: Table 4). Interestingly, this model showed a higher accuracy (precision and recall = 0.87) when tested on human DRS reads as compared to human cDNA reads (Additional file 1: Table 4). We also trained an MLP model (k=1,…,5) with the transcriptome annotation for *Candida glabrata* and tested it on *S. cerevisiae* ONT cDNA reads. This model yielded accuracy values as high as for the previous *S. cerevisiae* model (precision and recall = 0.94) (Fig. 2b) (Additional file 1: Table 4). As observed before for *S. cerevisiae* DRS reads, the model accuracy dropped when tested on DRS reads (precision and recall = 0.87) (Additional file 1: Table 4). We obtained similar results for the cross-species comparisons with the CNN model, with an improvement in accuracy for the mouse model applied to human DRS reads, and a drop for the *C. glabrata* model applied to *S. cerevisiae* DRS reads (Additional file 1: Table 4).

Reference-free interpretation of long-read transcriptome data generally involves some form of clustering (Marchet et al., 2018; Sahlin and Medvedev, 2018). Thus, to further demonstrate the utility of ReorientExpress for reference-free interrogation of transcriptomes with long-reads, we performed clustering of the cDNA reads (Methods). For the majority of clusters in human (>81%) and *S. cerevisiae* (>85%) ReorientExpress predicted correctly more than 50% of the reads in the cluster (Fig. 2c) (Additional file 1: Figure 1) (The proportion of clusters for each model can be found in Additional file 1: Table 5). That is, for most clusters, more than half the reads in those clusters can be correctly oriented. Accordingly, by taking the orientation of the cluster to be determined by that of the majority of reads, we could improve the overall orientation. To test this, we applied a majority vote per cluster to set the orientation of all reads in the cluster to be the majority label predicted by ReorientExpress. With this, ReorientExpress established the right orientation for the majority of cDNA reads for human and yeast, with up 96.2% of human reads and up to 98% of S. cerevisiae reads correctly oriented (Fig. 2d) (Additional file 1: Table 6).

### Comparisons with other models and inputs

Interestingly, inverting the procedure and training with ONT cDNA reads yields good accuracy when testing on annotated transcripts, but when training on ONT DRS reads the accuracy decreases (Additional file 1: Table 7). This could be a consequence of a higher proportion of base-calling errors in DRS reads due to the presence of RNA modifications, leading to a decrease in the identification of relevant sequence motifs learned by the model. To test this, we trained the MLP model with DRS reads from *in vitro* transcribed (IVT) RNA (Workman et al., 2018) and obtained slightly better accuracy than with DRS reads when testing on cDNA reads (Additional file 1: Table 7). Additionally, we observed no dependency with the basecaller used to obtain the sequence of reads. In particular, using Guppy-rapid or Guppy-high-accuracy to base-call the IVT RNA reads did not show any differences in the accuracy of the MLP model (Additional file 1: Table 8). This indicates that DRS errors may prevent accurate training of sequence-based models.

We also observed a dependency of the accuracy with the length of the reads. The prediction accuracy decreased for shorter reads (Additional file 1: Table 8), which suggests that either short molecules or partial reads may pose a limitation for the accurate prediction of orientation. To further test the effect of read length on the prediction accuracy, we trimmed a number of nucleotides from both ends of the cDNA reads in the test set. The accuracy was not significantly impacted performing trimming up to 200nt (Additional file 1: Table 9). Similarly, when we trimmed the training set by different amounts up to 200nt, leaving fixed the test set, the accuracy did not change significantly either (Additional file 1: Table 10). Thus, incomplete annotations can still be valid to train a model and complete annotations can yield accurate results on partial reads. This is relevant for the application to cDNA reads, which may be fragmented due to internal priming (Sessegolo et al., 2019). These results also indicate that DNN models are able to capture predictive features beyond the presence of adapters or poly-A tails to predict the 5’-to-3’ orientation of RNA molecules.

For comparison, we run pychopper (Methods), which can identify the orientation of cDNA reads by virtue of detecting the sequencing adapters. We analyzed all cDNA reads whose orientation was labelled previously. For the human cDNA reads, pychopper made predictions for only 24% of the reads, from which 98% were correctly classified. So in total, from the 270296 reads tested, ~23.5% (~63520 reads) were classified accurately by pychopper. This justifies the use of more sophisticated models to predict orientation. Additionally, we trained and tested a support vector machine (SVM) and a Random Forest (RF), using as inputs the same k-mer frequencies. Both also showed worse accuracy compared to the MLP model for the same test data. However, for *S. cerevisiae* the accuracy of both models trained with the *S. cerevisiae* annotation was high (Precision and recall 0.86 for the RF, and 0.95 for the SVM) (Additional File 1: Table 10). Finally, we also tested ReorientExpress with PacBio cDNA reads from Sorghum (Abdel-Ghany et al., 2016). We trained two MLP models, one with the Ensembl cDNA annotations from Sorghum (Sorghum bicolor NCBIv3) and another with Maize (Zea Mays B73_RefGen_v4). Both models showed high accuracy when tested against Sorghum PacBio reads (precision and recall ~0.95) (Additional file 1: Table 12).

### Association of RNA types and sequence motifs with read orientation prediction

Since we mapped long reads unambiguously to the transcriptome annotation for the purpose of benchmarking, we can use this information to investigate the accuracy of ReorientExpress according to the transcript type as provided by the Gencode annotation: protein-coding, processed transcript, lincRNA, etc. Applying the human CNN and MLP models (trained on 50000 random transcripts) to all labelled human cDNA reads, we observed that reads assigned to protein-coding transcripts, including transcripts from immunoglobulin related genes, showed the highest accuracy with more than 85% correctly predicted (Figs. 3a and 3b). Interestingly, even though transcript annotations of types “sense overlapping” and “sense intronic” were not used for training, reads assigned to them were correctly predicted in high proportion. In contrast, reads unambiguously associated with antisense or TEC (To be Experimentally Confirmed) transcripts, showed smaller accuracies compared with the rest of annotation types. TEC transcripts are based on EST clusters and may lack the sequence properties from other transcript types. Antisense transcripts remain difficult to predict correctly as they share sequence with transcripts annotated in the opposite strand. Nonetheless, inspection of the built clusters showed that in the majority of the clusters with antisense reads had all reads of type antisense. Indeed, 201 (78%) of clusters with at least one antisense read had 100% reads of type antisense, which corresponded to 882 antisense reads from the total of 976 antisense reads, i.e. 90% (Additional file 1: Figure 2). The CNN (Fig. 3a) and MLP (Fig. 3b) models showed very similar results, except for long non-coding RNAs (lncRNAs). Reads associated with lncRNAs from bidirectional promoters presented the same accuracy of the intergenic lncRNAs for the CNN model, but this was higher for the MLP model.

**Figure 3.**
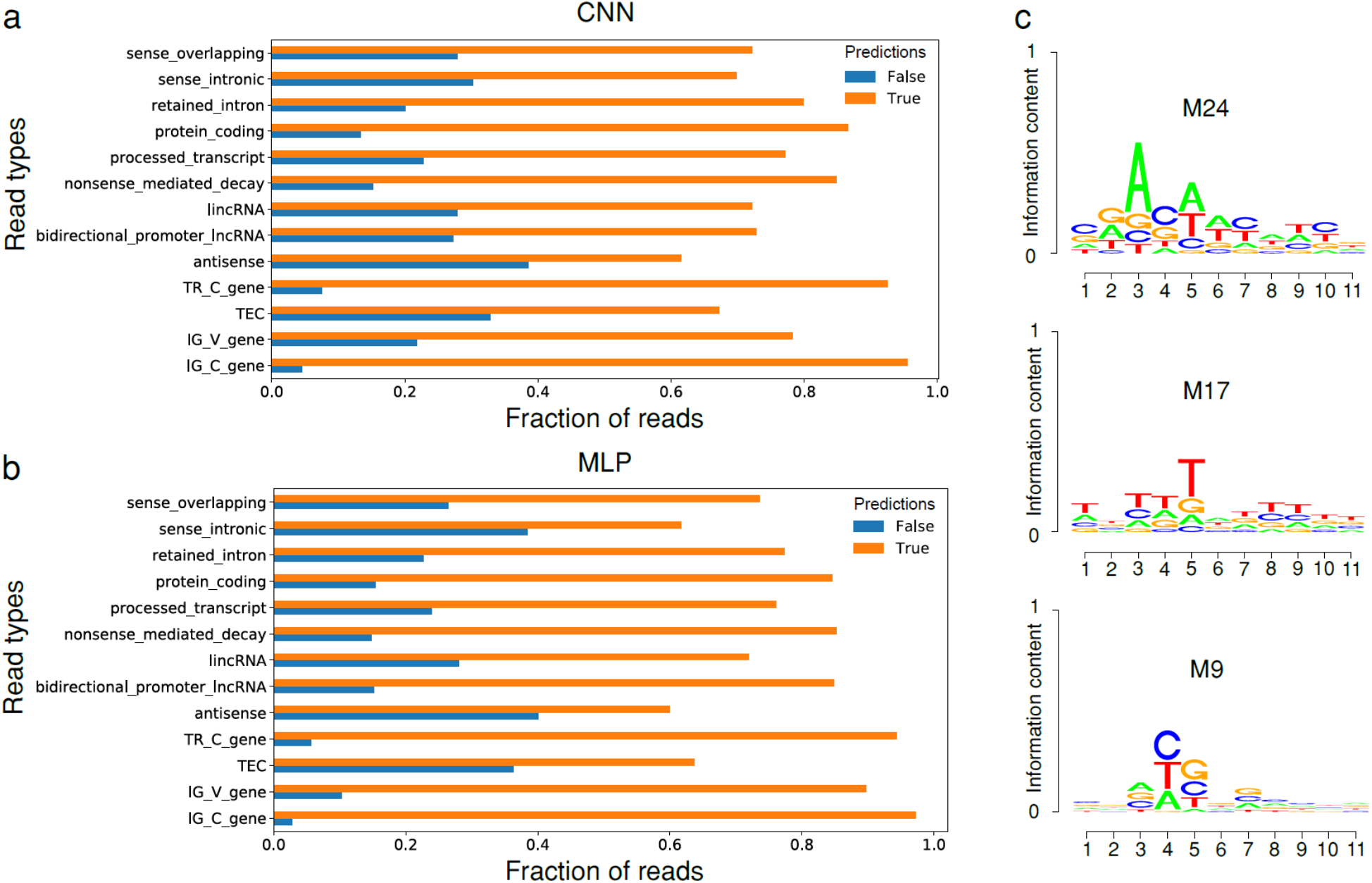
Read types and sequence motifs. The proportion of cDNA reads that were unambiguously mapped to each transcript type (y axis) and classified as correct (True) or incorrect (False) (x axis) by the CNN model **(a)** and the MLP model **(b)**. All transcript type annotations from the autosomes and sex chromosomes with more than 10 reads mapped are represented in the plot. **(c)** The motifs derived from three CNN filters with significant matches to previously described RNA-binding motifs (Additional file 3): filter M24 (RBM42, q-value = 0.0165997), filter M17 (HuR, q-value = 0.0312701), and filter M9 (PCBP1, q-value = 0.0426708). As information content (y axis) is low, the axis scale is given between 0 and 1.

To investigate whether ReorientExpress captures recognisable RNA motifs, we took advantage of the possibility to use the convolutional filters of the CNN to identify sequence motifs captured by the model as done previously (Alipanahi et al., 2015; Pan et al., 2018) (Methods). From these filters we found 32 candidate motifs (Additional file 2), which we compared with known protein-RNA binding motifs (Ray et al., 2013). This method detected motifs similar to those described for the RNA binding proteins PCBP1, ELAVL1 (HuR), and RBM42 (Fig 3b), among others (Additional file 3). Thus, sequence motifs that are relevant to predict molecule orientation recapitulate some of the binding specificities of proteins that control the metabolism of the RNA.

## Discussion

Here, we have shown that deep neural network (DNN) models trained on transcript sequences are able to provide an accurate orientation of cDNA long reads. We hypothesized that sequence motifs that are specific to RNA regulation would be identifiable in long sequencing reads despite the presence of errors, and found that some of the sequences relevant to predict molecule orientation are similar to known motifs involved in RNA-protein binding. The described DNN models maintain good accuracy despite using trimmed reads, and work well on nanopore and on PacBio reads. ReorientExpress provides a crucial aid in the interpretation of transcripts using cDNA long-reads in samples for which the genome reference is unavailable, as it is the case for many non-model organisms, but also in general for unstranded long-reads libraries for human and model organisms beyond the available references. This is particularly relevant considering the differences observed between individuals at large and small genomic scales (Dashnow et al., 2018; Sherman et al., 2019). Our analyses show that ReorientExpress can be very valuable in combination with long read clustering (Marchet et al., 2018; Sahlin and Medvedev, 2018) to facilitate more accurate downstream analyses of transcriptomes, like the prediction of open reading frames. The ability to predict the 5’-to-3’ orientation of cDNA long reads using models trained on related species, makes ReorientExpress a key processing tool for the study of transcriptomes from non-model organisms with long-reads.

## Methods

### Training and testing ReorientExpress

ReorientExpress (https://github.com/comprna/reorientexpress) implements deep neural network (DNN) models in keras (https://github.com/keras-team/keras) and Tensorflow (Abadi et al., 2016). All input data is preprocessed to discard reads that contain N’s. For reads from direct RNA-seq experiments, uracil (U) is transformed into thyimine (T). Input reads can be optionally trimmed and this is done for the same length on both sides of each input sequence. For training purposes, a random selection of half the sequences are reverse-complemented to obtain a balanced training set. Optionally, all sequences can be reverse complemented to double up the training input. ReorientExpress implements two different DNN models, a multi-layer perceptron (MLP) and a convolutional neural network (CNN). In the MLP model, sequences are processed to build a matrix of k-mer frequencies, from k=1 up to a specified k-mer length (default k=1,...,5). The normalization is performed per input sequence and per k-mer length. That is, for a fixed k, each k-mer count is divided by the total number of k-mers in the sequence of length L, so that frequency(k-mer) = count(k-mer)/(L-k+1). Using the k-mer frequencies ensures that the input size is the same for all transcripts regardless of the transcript length. MLPs are simpler than Convolutional (CNNs), so they are faster to train and to run. On the other hand, CNNs can model relative spatial relationships, hence they can take sequence context into account. For this reason, we also included a CNN model in ReorientExpress. For the CNN model, each input sequence was divided into overlapping sequences of 500nt, overlapping by 250nt. For transcripts of length between 250 and 500 we added Ns at the end of the sequence. We used one hot encoding as input for each one of the 500nt windows.

Once a model is trained, or given an already available model, ReorientExpress can predict the orientation of a set of unlabeled reads in *prediction* mode. ReorientExpress feeds the normalized k-mer counts for each read for the MLP model, or the sliding windows for the CNN model to predict the orientation. In the MLP model, the last layer has only one node, which applies a sigmoid function to approximate a probability from the score it receives. The probability can be interpreted as the certainty that the input read is not in the correct orientation. So, a read with a score greater than 0.5 is predicted to be in the wrong orientation and is reverse-complemented. For the CNN model, for each window tested the output is a posterior of the orientation given that window. To provide a prediction for each input read, ReorientExpress takes the mean value for both orientations independently, and outputs the orientation with the greatest mean.

The *test* mode is aimed at evaluating the accuracy of a model using as input sequences with known orientation. The program generates predictions for the input reads and compares them with the provided labels, returning a precision (proportion of the predictions that are correct), a recall (true positive rate, proportion of labeled cases that are correctly predicted), an F1-score (harmonic mean of precision and recall) and the total number of input reads. As input for any of the three modes, *train*, *predict* and *test*, one can use three types of datasets: *experimental*, *annotation* or *mapped*. Experimental data refers to any kind of long-read data for which the orientation is known, such as direct RNA-seq, and reads are considered to be given in the 5’-to-3’ orientation. Annotation data refers to the transcript sequences from a reference annotation, such as the human transcriptome reference. Annotation is considered to be in the right 5’-to-3’ orientation, and can include the transcript type, such as protein coding, processed transcript, etc. Mapped data refers to sequencing data, usually cDNA, whose orientation has been annotated by an independent method, e.g. by mapping the reads to a reference. In this case, a PAF file for the mapping, together with the FASTA/FASTQ file, is required. The labelled data is used for training or testing. In *predict* mode the data does not require labelling and ReorientExpress provides a prediction. More details are provided in https://github.com/comprna/reorientexpress.

### Deep Neural Network (DNN) models tested

Models used for the analyses described in the manuscript are provided at https://github.com/comprna/reorientexpress. The human model was trained using the Gencode annotation release 28, and the mouse model was built using the mouse Gencode release M19. The Ensembl annotation (https://fungi.ensembl.org/) was used to train the *Saccharomyces cerevisiae* (R64-1-1) and the *Candida glabrata* (ASM254v2) models. Ensembl annotations (http://plants.ensembl.org) were used from Sorghum (Sorghum bicolor NCBIv3) and from Maize (Zea Mays B73_RefGen_v4) to build models to test on PacBio data. From the annotation files, we only used the most frequent transcript annotation types: protein coding, lincRNA, processed transcripts, antisense and retained intron. We trained the models using 50,000 randomly selected transcript sequences from the annotation, or all of them if there were less than 50,000 (*S. cerevisiae* and C. *glabrata*). The results did not change when running the analysis with different sets of 50,000 transcripts.

### Test datasets

To test ReorientExpress on cDNA reads, we first calculated a set of cDNA reads for which orientation could be determined unambiguously in an independent way. We used human cDNA from the Nanopore consortium (cDNA 1D pass reads from JHU run 1) (Workman et al., 2018) (available from https://github.com/nanopore-wgs-consortium/NA12878/blob/master/nanopore-human-transcriptome/fastq_fast5_bulk.md) and *S. cerevisiae* cDNA reads (Garalde et al., 2018) from SRA (SRR6059708). We mapped the cDNA reads to the corresponding transcriptome annotation using minimap2 (Li, 2018) without secondary alignments (minimap2 -cx map-ont -t7 --secondary=no). We kept only reads with maximum mapping quality (MAPQ = 60) and that were uniquely mapping. For human, 899431 out of 962598 reads were mapped in this way, 282444 of which had MAPQ = 60. After removing the ~4% multimapping cases, we finally obtained 270296 reads with orientation unambiguously assigned. For *S. cerevisiae*, 4000698 out of a total of 5045243 reads were mapped, 3089543 of which had MAPQ = 60. After removing the ~3% multimapping cases, we finally obtained 2984873 reads with orientation unambiguously assigned. Additionally, we used direct RNA sequencing (DRS) for human (JHU Run 1 available from https://github.com/nanopore-wgs-consortium/NA12878/blob/master/nanopore-human-transcriptome/fastq_fast5_bulk.md) and for *S. cerevisiae* from SRA (SRR6059706) (Garalde et al., 2018). We also tested ReorientExpress with PacBio cDNA reads from Sorghum (Abdel-Ghany et al., 2016) (Data available at https://zenodo.org/record/49944#.XCkXQC-ZN24). We trained two MLP models with the Ensembl cDNA annotations (http://plants.ensembl.org) from Sorghum (Sorghum bicolor NCBIv3) and Maize (Zea Mays B73_RefGen_v4).

### Other models for comparison

We tested a Random Forest model and an SVM model using sklearn libraries. The models were trained using 50,000 random annotated transcripts, using as training input the normalized k-mer frequencies for each sequence as features and the orientation as classification label. Further details are provided in Additional file 1. We also run pychopper (https://github.com/nanoporetech/pychopper) (cdna_classifier.py command with the list of barcodes provided by pychopper) on the 270296 human cDNA reads that we had labelled previously.

### Testing the dependency with base-callers

We used Guppy rapid and Guppy high accuracy (v2.2.3) with the signal files from the in-vitro transcript RNA sequenced with MinION by the Nanopore Consortium (available from https://github.com/nanopore-wgs-consortium/NA12878/blob/master/nanopore-human-transcriptome/fastq_fast5_bulk.md). As this is direct RNA sequencing, the orientation of the reads can be readily used to test the accuracy of our models.

### Clustering and majority vote

We performed clustering of the human and *S. cerevisiae* cDNA reads using IsONclust (Sahlin and Medvedev, 2018). Only cDNA reads that had been assigned an orientation by mapping as described above were used for clustering. We predicted the read 5’-to-3’ orientation for the same reads with ReorientExpress and calculated for each cluster the proportion of reads that were correctly orientated. As IsONclust does not given clusters with oriented reads, the orientation of all cDNA reads was taken from the mapping described above. In each cluster we then predicted the read orientation with ReorientExpress and selected the majority label to assign all reads in the cluster: if the majority (>50%) of reads were predicted to be already in 5’-to-3’ orientation (forward), we set all reads to forward. Otherwise, all reads were reverse-complemented. The accuracy of all reads was then calculated by comparing our predictions with the predetermined orientations.

### Motif analysis

We studied the 32 filters from the first layer of the CNN to obtain the sequences that are most informative for predicting the orientation, using an approach similar to (Alipanahi et al., 2015; Pan et al., 2018). To explore exhaustively all potential motifs, we used activations above 0 and converted the associated sequences to position weight matrices (PWMs). The derived 32 motif matrices (Additional file 2) were then compared against the CISBP-RNA database (http://cisbp-rna.ccbr.utoronto.ca/) (Ray et al., 2013) using the TOMTOM algorithm (http://meme-suite.org/doc/tomtom.html) (Gupta et al., 2007) for the comparison of PWM-based motifs and selecting matches with p−value < 0.05 (Additional file 3).

## List of Abbreviations

ONT: Oxford Nanopore Technologies
DRS: Direct RNA sequencing
cDNA: complementary DNA
k-mer: length k oligomer
DNN: Deep Neural Network
MLP: Multi-layer perceptron
CNN: Convolutional Neural Network
IVT: In vitro transcribed
PWM: Position Weight Matrix

## Competing interests

None of the authors have any competing interests.

## Availability of data and materials

All datasets on which the conclusions of the paper rely are accessible from publicly available repositories and referenced accordingly. The code generated for this article is available at https://github.com/comprna/reorientexpress

**Additional file 1:** Additional figures and tables referenced in the article.

**Additional file 2:** Motif file for the 32 filters from the CNN model.

**Additional file 3:** Matches of the 32 motifs from Additional file 2 to RBP motifs from CISBP-RNA database.

## Notes

#### Summary of Updates

We provide extra analyses and information about the methods used. We have included a convolutional neural network for comparison. We have also described how the accuracy depends on the transcript type the read originates from. Additionally, we have characterized the sequence motifs that contribute to the prediction of the read orientation, and have identified similarities with motifs for RNA-protein interactions.

https://github.com/comprna/reorientexpress

