## Additional file 1 for "ReorientExpress: Reference-free orientation of nanopore cDNA reads with deep learning"

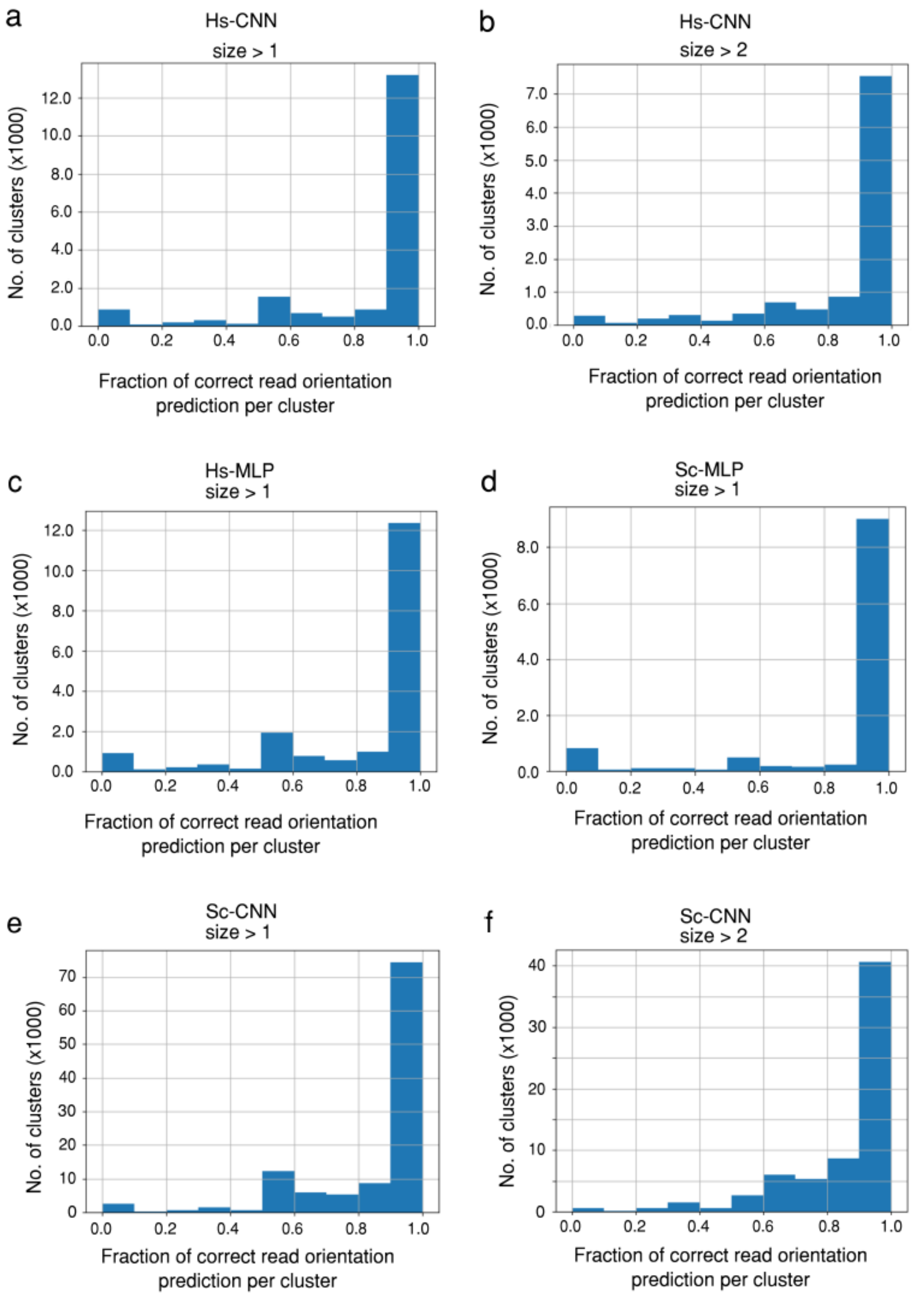

**Figure 1. Proportion of corrected reads per cluster.** Number of clusters (y axis) according to the proportion of ONT cDNA reads in the cluster with orientation correctly predicted by ReorientExpress (x axis) for the CNN model trained on the human transcriptome (Hs-CNN) for clusters with >1 reads **(a)** and for clusters with >2 reads **(b)**, for the MLP model trained on the human transcriptome (Hs-CNN) for clusters with >1 reads **(c)**, for the MLP model trained on the S. cerevisiae transcript (Sc-MLP) for clusters with >1 reads **(d)**, and for the CNN model trained on the S. cerevisiae transcriptome (Sc-CNN) for clusters with >1 reads **(e)** and for clusters with >2 reads **(f).** The number of clusters with more than 50% of their reads correctly oriented are given in Table 5. The total number of reads correctly oriented after using a majority vote are given in Table 6.

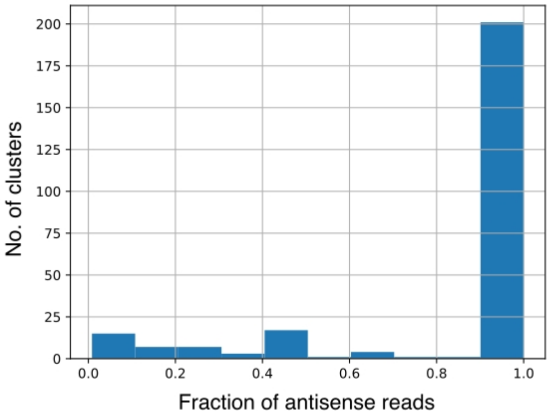

**Figure 2. Clusters with antisense cDNA reads.** The plot shows the clusters that contained 1 or more reads labelled as antisense, according to the proportion of reads in each cluster that were of type antisense. The plot shows that the majority of clusters with antisense reads consists of 100% antisense reads. The clusters that have 100% of their reads labeled as antisense are 201 (78%), which correspond to 882 antisense reads from the total of 976 antisense reads, i.e. 90%.

| Layer | Type | Activation | Nodes | Dropout Rate |
|---|---|---|---|---|
| 0 | Input | - | 1364 | |
| 1 | Dense | ReLu | 500 | |
| | Dropout | - | | 0.3 |
| 2 | Dense | ReLu | 250 | |
| | Dropout | - | | 0.3 |
| 3 | Dense | ReLu | 125 | |
| | Dropout | - | | 0.3 |
| 4 | Dense | ReLu | 62 | |
| | Dropout | - | | 0.3 |
| 5 | Dense | ReLu | 31 | |
| | Dropout | - | | 0.3 |
| 6 | Dense | Sigmoid | 1 | |

**Table 1. Architecture of the MLP model.** First layer is the input, with as many nodes as the number of normalized frequencies. For k-mers with k=1,...,5, there are 4 + 16 + 64 + 256 + 1024 = 1364 frequencies. For every layer (column Layer), the activation function (Activation) and the number of nodes (Nodes) is described. The activation functions used are the Rectified Linear Unit (ReLu) and the sigmoid. For each layer, the Dropout layer inserted immediately after the dense layer and the dropout rate is indicated.

| Layer | Type | Activation | No. of filters | Filter size/nodes | Dropout Rate |
|---|---|---|---|---|---|
| 1 | Conv1 | ReLu | 32 | 11 x 4 | - |
| | Pool1 | - | 32 | 4 x1 | - |
| 2 | Conv2 | ReLu | 64 | 3 x 1 | - |
| | Pool2 | - | 64 | 2 x1 | - |
| 3 | Conv3 | ReLu | 96 | 3 x 1 | - |
| | Pool3 | - | 96 | 2 x1 | - |
| 4 | Flatten | - | - | - | - |
| 5 | Dense | ReLu | - | 256 | - |
| | Dropout | - | - | - | 0.4 |
| 6 | Dense | ReLu | - | 256 | - |
| 7 | Dense | softmax | - | 2 | - |

**Table 2. Architecture of the CNN model.** The model consists of 3 convolutional layers (Conv1-3), 3 pooling layers (Pool1-3) and 3 dense layers, with different filter sizes. The Conv and Pool layers identify the important features and the dense layers combine these features together to make the prediction. The activation function used are the Rectified Linear Unit (ReLu) and the softmax. The dropout of 0.4 is used between first and second dense layer.

| | | | Forward | | | | Reverse | | | | Average | | | |
|---|---|---|---|---|---|---|---|---|---|---|---|---|---|---|
| Model | Training | Testing | Prec. | Recall | F1 | Reads | Prec. | Recall | F1 | Reads | Prec. | Recall | F1 | Reads |
| MLP | Human transcriptome | human cDNA reads | 0.76 | 0.84 | 0.80 | 20655 | 0.89 | 0.82 | 0.85 | 30450 | 0.84 | 0.83 | 0.83 | 51105 |
| MLP | Human transcriptome | Human DRS reads | 0.86 | 0.86 | 0.86 | 25081 | 0.86 | 0.86 | 0.86 | 24919 | 0.86 | 0.86 | 0.86 | 50000 |
| MLP | S. cerevisiae transcriptome | S. cerevisiae cDNA reads | 0.91 | 0.93 | 0.92 | 19962 | 0.95 | 0.94 | 0.94 | 30460 | 0.93 | 0.93 | 0.93 | 50422 |
| MLP | S. cerevisiae transcriptome | S. cerevisiae DRS reads | 0.85 | 0.94 | 0.89 | 25023 | 0.93 | 0.83 | 0.88 | 24977 | 0.89 | 0.88 | 0.88 | 50000 |
| CNN | Human transcriptome | human cDNA reads | 0.79 | 0.90 | 0.84 | 21476 | 0.92 | 0.82 | 0.86 | 28525 | 0.86 | 0.86 | 0.85 | 50001 |
| CNN | Human transcriptome | Human DRS reads | 0.91 | 0.91 | 0.91 | 50000 | 0.91 | 0.91 | 0.91 | 50000 | 0.91 | 0.91 | 0.91 | 50000 |
| CNN | S. cerevisiae transcriptome | S. cerevisiae cDNA reads | 0.86 | 0.91 | 0.88 | 23095 | 0.92 | 0.87 | 0.90 | 26906 | 0.89 | 0.89 | 0.89 | 50001 |
| CNN | S. cerevisiae transcriptome | S. cerevisiae DRS reads | 0.77 | 0.88 | 0.82 | 25000 | 0.86 | 0.74 | 0.79 | 25000 | 0.82 | 0.81 | 0.80 | 50000 |

**Table 3. DNN models trained on a transcriptome and tested on cDNA or direct RNA reads from the same species.** Precision, recall (true positive rate), and F1-score for the models are given for each orientation separately, and for the total set. The number of reads tested are also provided. The human cDNA (1D) and DRS data corresponds to the JHU Run 1 available from https://github.com/nanopore-wgs-consortium/NA12878/blob/master/nanopore-human-transcriptome/fastq_fast5_bulk.md). The S. cerevisiae nanopore cDNA (SRR6059708) and DRS (SRR6059706) reads were obtained from SRA (SRP118556) (Garalde et al., 2018).

| | | | Forward | | | | Reverse | | | | Average | | | |
|---|---|---|---|---|---|---|---|---|---|---|---|---|---|---|
| Model | Training | Testing | Prec. | Recall | F1 | Reads | Prec. | Recall | F1 | Reads | Prec. | Recall | F1 | Reads |
| MLP | mouse transcriptome | human cDNA reads | 0.60 | 0.94 | 0.73 | 21255 | 0.92 | 0.55 | 0.69 | 28745 | 0.79 | 0.71 | 0.71 | 50000 |
| MLP | mouse transcriptome | Human DRS reads | 0.88 | 0.86 | 0.87 | 24327 | 0.86 | 0.88 | 0.87 | 24326 | 0.87 | 0.87 | 0.87 | 48653 |
| MLP | C. glabrata transcriptome | S. cerevisiae cDNA reads | 0.96 | 0.93 | 0.94 | 23486 | 0.94 | 0.97 | 0.95 | 26514 | 0.95 | 0.95 | 0.95 | 50000 |
| MLP | C. glabrata transcriptome | S. cerevisiae DRS reads | 0.85 | 0.90 | 0.87 | 24844 | 0.90 | 0.84 | 0.87 | 24843 | 0.87 | 0.87 | 0.87 | 49687 |
| MLP | Human transcriptome | S. cerevisiae DRS reads | 0.77 | 0.79 | 0.78 | 24844 | 0.78 | 0.77 | 0.78 | 24843 | 0.78 | 0.78 | 0.78 | 49687 |
| MLP | Human transcriptome | Sorghum PacBio reads | 0.61 | 0.61 | 0.61 | 37239 | 0.61 | 0.62 | 0.62 | 37239 | 0.62 | 0.62 | 0.62 | 74478 |
| CNN | Mouse transcriptome | human cDNA reads | 0.79 | 0.90 | 0.84 | 21476 | 0.92 | 0.82 | 0.87 | 28525 | 0.85 | 0.86 | 0.85 | 50001 |
| CNN | Mouse transcriptome | Human DRS reads | 0.90 | 0.93 | 0.92 | 25000 | 0.93 | 0.90 | 0.91 | 25000 | 0.92 | 0.92 | 0.92 | 50000 |
| CNN | C. glabrata transcriptome | S. cerevisiae cDNA reads | 0.94 | 0.93 | 0.94 | 23095 | 0.94 | 0.95 | 0.95 | 26906 | 0.94 | 0.94 | 0.94 | 50001 |
| CNN | C. glabrata transcriptome | S. cerevisiae DRS reads | 0.86 | 0.89 | 0.88 | 25000 | 0.89 | 0.86 | 0.87 | 25000 | 0.88 | 0.88 | 0.88 | 50000 |

**Table 4. Cross-species DNN models.** Precision, recall (true positive rate), and F1-score for the model trained on the annotated transcriptome for mouse tested on human nanopore cDNA (1D JHU run 1) and direct RNA sequencing (DRS) reads (JHU, run 1) reads from (Workman et al., 2018) (https://github.com/nanopore-wgs-consortium/NA12878/blob/master/RNA.md). Orientation labels for cDNA reads were previously calculated by mapping them to the transcriptome (see Methods). The accuracy values for the reads oriented as obtained after mapping, and reverse-complemented from the mapping (Reverse), and the average value is provided. The number of reads used for testing is shown in the corresponding columns.. *S. cerevisiae* nanopore direct RNA reads were obtained from (Garalde et al., 2018) (https://trace.ncbi.nlm.nih.gov/Traces/sra/?study=SRP118556) and PacBio cDNA reads for Sorghum were obtained from (Abdel-Ghany et al., 2016) (https://zenodo.org/record/49944#.XCkXQC-ZN24 ).

| Model | Cluster with >1 read | Clusters with >2 reads |
|---|---|---|
| Hs_MLP | 81.21 % | 87.68 % |
| Hs_CNN | 83.91 % | 89.29 % |
| Sc_MLP | 85.82 % | 88.11 % |
| Sc_CNN | 85.43 % | 92.85 % |

**Table 5. Clusters with more than 50% of reads correctly oriented.** The table shows the percentage of clusters having more than 50% of the reads correctly oriented by ReorientExpress separated by species, human (Hs) or *S. cerevisiae* (Sc), and DNN model, MLP or CNN. The results are shown for clusters with more than 1 read or with more than 2 reads.

| **Proportions** | **Hs_CNN** | **Hs_MLP** | **Sc_CNN** | **Sc_MLP** |
|---|---|---|---|---|
| **Default** | 42.51 % | 42.51 % | 47.12 % | 47.12 % |
| **ReorientExpress** | 85.44 % | 83.77 % | 88.68 % | 86.75 % |
| **ReorientExpress and clustering** | 96.20 % | 95.56 % | 98.05 % | 90.24 % |
| **Total Number of reads** | **Hs_CNN** | **Hs_MLP** | **Sc_CNN** | **Sc_MLP** |
| **Default** | 114902 | 114902 | 1406529 | 1406529 |
| **ReorientExpress** | 230942 | 226425 | 2646930 | 2589360 |
| **ReorientExpress and clustering** | 260033 | 258289 | 2926643 | 2693432 |

**Table 6. Reads correctly oriented.** Proportions (upper table) and total number of reads correctly oriented from the Human (Hs) cDNA sample (270296 labelled reads) and the S. cerevisiae (Sc) cDNA sample (2984873 labelled reads).

| | | | Forward | | | | Reverse | | | | Average | | | |
|---|---|---|---|---|---|---|---|---|---|---|---|---|---|---|
| Model | Training | Testing | Prec. | Recall | F1 | Reads | Prec. | Recall | F1 | Reads | Prec. | Recall | F1 | Total Reads |
| MLP | Human cDNA reads | human transcriptome | 0.85 | 0.81 | 0.83 | 25000 | 0.85 | 0.81 | 0.83 | 25000 | 0.83 | 0.83 | 0.83 | 50000 |
| MLP | Human DRS reads | Human cDNA reads | 0.67 | 0.70 | 0.68 | 20655 | 0.79 | 0.76 | 0.77 | 30450 | 0.74 | 0.74 | 0.74 | 51105 |
| MLP | Human IVT RNA DRS reads | Human cDNA reads | 0.75 | 0.61 | 0.67 | 21690 | 0.74 | 0.84 | 0.79 | 28320 | 0.74 | 0.72 | 0.73 | 50001 |
| CNN | Human IVT RNA DRS reads | Human cDNA reads | 0.80 | 0.72 | 0.75 | 21476 | 0.80 | 0.86 | 0.83 | 28525 | 0.80 | 0.79 | 0.79 | 50001 |
| MLP | S. cerevisiae cDNA reads | S. cerevisiae transcriptome | 0.87 | 0.88 | 0.88 | 3299 | 0.88 | 0.88 | 0.88 | 3299 | 0.88 | 0.88 | 0.88 | 6598 |

**Table 7. Models trained on Nanopore reads.** Precision (Prec.), recall (true positive rate), and F1-score for the model trained on cDNA or direct RNA nanopore reads from (Workman et al., 2018) (https://github.com/nanopore-wgs-consortium/NA12878/blob/master/RNA.md) for human, and tested on the human transcriptome; or trained on S. cerevisiae cDNA reads and tested on the *S. cerevisiae* transcriptome. The accuracy values for the reads oriented as obtained after mapping, and reverse-complemented from the mapping (Reverse), and the average value is provided. The number of reads used for testing is shown in the corresponding columns.

| | Guppy - fast algorithm | Guppy - high accuracy |
|---|---|---|
| **Read length (bp)** | **% correctly predicted** | **% correctly predicted** |
| >50 | 54% | 54% |
| >250 | 86% | 87% |
| >500 | 92% | 92% |

**Table 8. Testing the dependency with base-calling.** Guppy fast and Guppy high accuracy with the signal files from the in-vitro transcript RNA sequenced with MinION by the Nanopore Consortium (available from https://github.com/nanopore-wgs-consortium/NA12878/blob/master/nanopore-human-transcriptome/fastq_fast5_bulk.md ) are used. As this is direct RNA sequencing, the orientation of the reads can be used to test the accuracy of our models. The test was performed using 50,000 random IVT RNA reads with the MLP model trained on 50,000 random transcripts from human.

| | Forward | | | | Reverse | | | | Average | | | |
|---|---|---|---|---|---|---|---|---|---|---|---|---|
| Trim | Precision | Recall | F1-score | Reads | Precision | Recall | F1-score | Reads | Precision | Recall | F1-score | Total reads |
| 10nt | 0.78 | 0.83 | 0.80 | 8468 | 0.88 | 0.85 | 0.86 | 12755 | 0.84 | 0.84 | 0.84 | 21223 |
| 20nt | 0.78 | 0.83 | 0.80 | 8528 | 0.88 | 0.84 | 0.86 | 12892 | 0.84 | 0.84 | 0.84 | 21420 |
| 40nt | 0.77 | 0.82 | 0.80 | 8531 | 0.88 | 0.84 | 0.86 | 12916 | 0.83 | 0.83 | 0.83 | 21447 |
| 80nt | 0.76 | 0.82 | 0.79 | 8518 | 0.88 | 0.83 | 0.85 | 12855 | 0.83 | 0.83 | 0.83 | 21373 |
| 100nt | 0.75 | 0.82 | 0.78 | 8521 | 0.86 | 0.83 | 0.84 | 12957 | 0.81 | 0.82 | 0.82 | 21478 |
| 200nt | 0.74 | 0.80 | 0.77 | 8418 | 0.82 | 0.80 | 0.81 | 12825 | 0.79 | 0.80 | 0.79 | 21253 |

**Table 9. Testing on trimmed reads.** The human model (trained on the transcriptome) and tested on the human cDNA reads after trimming a certain number of nucleotides from both ends of the read: 10, 20, 40, 80, 100, and 200. In each case, the accuracy values are calculated as before.

| | Forward | | | | Reverse | | | | Average | | | |
|---|---|---|---|---|---|---|---|---|---|---|---|---|
| Trim | Precision | Recall | F1-score | Reads | Precision | Recall | F1-score | Reads | Precision | Recall | F1-score | Reads |
| 0 | 0.71 | 0.81 | 0.76 | 5139 | 0.84 | 0.76 | 0.80 | 6898 | 0.79 | 0.78 | 0.78 | 12037 |
| 50 | 0.73 | 0.80 | 0.76 | 5136 | 0.84 | 0.78 | 0.81 | 7158 | 0.79 | 0.79 | 0.79 | 12474 |
| 100 | 0.72 | 0.79 | 0.75 | 5359 | 0.83 | 0.78 | 0.80 | 7241 | 0.79 | 0.78 | 0.78 | 12600 |
| 150 | 0.69 | 0.79 | 0.74 | 5329 | 0.82 | 0.74 | 0.78 | 7184 | 0.77 | 0.76 | 0.76 | 12513 |
| 200 | 0.67 | 0.81 | 0.73 | 5170 | 0.83 | 0.70 | 0.76 | 6924 | 0.76 | 0.75 | 0.75 | 12094 |

**Table 10. Training with trimmed transcripts.** The MLP human models were trained on the transcriptome after trimming the corresponding nucleotides from both ends. The test was performed on the human cDNA reads without any trimming.

| | | | Forward | | | | Reverse | | | | Average | | | |
|---|---|---|---|---|---|---|---|---|---|---|---|---|---|---|
| Model | Training | Testing | Prec. | Recall | F1 | Reads | Prec. | Recall | F1 | Reads | Prec. | Recall | F1 | Total Reads |
| Random Forest | Human transcriptome | human cDNA reads | 0.65 | 0.69 | 0.67 | 21255 | 0.76 | 0.72 | 0.74 | 28745 | 0.71 | 0.71 | 0.71 | 50000 |
| Random Forest | S. cerevisiae transcriptome | S. cerevisiae cDNA reads | 0.87 | 0.82 | 0.85 | 23486 | 0.85 | 0.89 | 0.87 | 26514 | 0.86 | 0.86 | 0.86 | 50000 |
| SVM | Human transcriptome | human cDNA reads | 0.71 | 0.69 | 0.70 | 21255 | 0.78 | 0.79 | 0.78 | 28745 | 0.75 | 0.75 | 0.75 | 50000 |
| SVM | S. cerevisiae transcriptome | S. cerevisiae cDNA reads | 0.96 | 0.93 | 0.95 | 23486 | 0.94 | 0.97 | 0.95 | 28745 | 0.95 | 0.95 | 0.95 | 50000 |

**Table 11. Accuracy of Random Forest and SVM models.** Precision (Prec.), recall (true positive rate), and F1-score (F1) for each model based on k-mers k=1,…,5, trained on the annotated human or S. cerevisiae transcriptomes and tested on nanopore cDNA reads from the same species (JHU, 1D cDNA reads, run 1, from NA12878 for human https://github.com/nanopore-wgs-consortium/NA12878/blob/master/nanopore-human-transcriptome/fastq_fast5_bulk.md, and SRR6059708 for S. cerevisiae). For the Random Forest the default parameters were used except for the maximum depth, which was tested from 1 to 10. The best accuracy was obtained with depth 9, which is the one reported. The complete list of parameters used for training the random forest and SVM is provided below.

*RandomForestClassifier(bootstrap=True, class_weight=None, criterion='gini', max_depth=9, max_features='auto', max_leaf_nodes=None, min_impurity_decrease=0.0, min_impurity_split=None, min_samples_leaf=1, min_samples_split=2, min_weight_fraction_leaf=0.0, n_estimators=10, n_jobs=1, oob_score=False, random_state=0, verbose=0, warm_start=False).*

For the SVM different kernels: rbf, linear and poly were tested. The best accuracy model parameters are below:

*SVC(C=1.0, cache_size=200, class_weight=None, coef0=0.0, decision_function_shape='ovr', degree=3, gamma='auto', kernel='linear', max_iter=-1, probability=False, random_state=None, shrinking=True, tol=0.001, verbose=False)*.

Incidentally, the SVM was much slower than RFs. For comparison: for the human dataset and using the same computer, the RF took 421 seconds (around 7 min) but the SVM took 47825 seconds (>13 hours), i.e. 100 times slower.

| | Forward | | | | Reverse | | | | Average | | | |
|---|---|---|---|---|---|---|---|---|---|---|---|---|
| | Prec. | Recall | F1 | Reads | Prec. | Recall | F1 | Reads | Prec. | Recall | F1 | Reads |
| Sorghum model tested on Sorghum Pacbio | 0.95 | 0.95 | 0.95 | 25000 | 0.95 | 0.95 | 0.95 | 25000 | 0.95 | 0.95 | 0.95 | 50000 |
| Maize model tested on Sorghum Pacbio | 0.94 | 0.95 | 0.95 | 25000 | 0.95 | 0.94 | 0.95 | 25000 | 0.95 | 0.95 | 0.95 | 50000 |

**Table 12. Testing of ReorientExpress with PacBio data.** Precision, recall (true positive rate), and F1-score for the model trained on the annotated transcriptome for Sorghum (Sorghum bicolor NCBIv3) and Maize (Zea Mays B73_RefGen_v4) tested on PacBio cDNA reads for Sorghum (Abdel-Ghany et al., 2016) (https://zenodo.org/record/49944#.XCkXQC-ZN24). The accuracy values for the reads oriented as obtained after mapping, and reverse-complemented from the FASTA/FASTQ, and the average value is provided. The number of reads used for testing is shown in the corresponding columns.
